# Familiarity-taxis: A bilateral approach to view-based navigation

**DOI:** 10.1101/2023.06.23.546247

**Authors:** Fabian Steinbeck, Efsthathios Kagioulis, Alex Dewar, Andy Philippides, Thomas Nowotny, Paul Graham

## Abstract

Many insects use view-based navigation, or snapshot matching, to return to familiar locations, or navigate routes. This relies on egocentric memories being matched to current views of the world. Previous snapshot navigation algorithms have used full panoramic vision for the comparison of memorised images with query images to establish a measure of familiarity, which leads to a recovery of the original heading direction from when the snapshot was taken. Many aspects of insect sensory systems are lateralised with steering being derived from the comparison of left and right signals like a classic Braitenberg vehicle. Here we investigate whether view-based route navigation can be implemented using bilateral visual familiarity comparisons. We found that the difference in familiarity between estimates from left and right fields of view can be used as a steering signal to recover the original heading direction. This finding extends across many different sizes of field of view and visual resolutions. In insects, steering computations are implemented in a brain region called the Lateral Accessory Lobe, within the Central Complex. In a simple simulation we show with a SNN model of the LAL an existence proof of how bilateral visual familiarity could drive a search for a visually defined goal.

## Introduction

Visual navigation is essential for many animals and has increasing usage in artificial systems, as it presents a cheap way to provide egocentric spatial information for application in autonomous agents. Effective visual navigation algorithms enable autonomous agents to explore and navigate environments without external control. In contrast to methods such as Simultaneous Localization and Mapping method (SLAM, Smith & Cheeseman (1986)), insect inspired view-based navigation models derive from agents with poor visual resolution (1°-4°, Schwarz et al. (2011)) and computationally restricted hardware (ants have fewer than *∼* 500,000 neurons in their brain, (Godfrey et al., 2021). Thus, mimicking insect behaviours and computational principles may overcome hardware limitations and increase not only computational efficiency, but, given the navigational prowess of insects, also help with the robustness of applied navigation mechanisms in dynamic environments.

Behavioural experiments and modelling of visual landmark learning in bees (Cartwright & Collett, 1983), sparked the subsequent development of several view-based navigation algorithms (Franz et al., 1998; Chahl & Srinivasan, 1996; Möller & Vardy, 2006; Smith et al., 2007). With view-based navigation, an environment can be sparsely represented with reference images (snapshots) taken at nodes relevant for navigation within the environment. For instance, navigation to a node can happen via gradient descent, as the image difference can be used as an indicator for proximity to a goal location (Zeil et al., 2003). By calculating the root mean square pixel difference between panoramic snapshots and query images from displaced locations in the vicinity, a spatial Image Difference Function (IDF) can be established. This function is minimal when the query image is in the same location as the snapshot, but gradually increases with distance to that snapshot’s location. By following the descending gradient, the location of the snapshot can be found.

The measure of visual difference can also be directly incorporated into an algorithm for setting orientation as part of homing algorithms. One set of methods uses the concept of a ‘visual compass’ (Labrosse, 2006; Zeil et al., 2003) which is a method to find the best matching orientation between a snapshot and a query image, therefore leading to the recovery of the heading at which the snapshot was stored. Systematic rotational sampling, in order to find the best match with a stored shapshot, resembles the behaviour of ants which scan the environment with body rotations while visually navigating (Lent et al., 2010; Wystrach et al., 2014). Such visual compass algorithms are good at explaining the route navigation behaviours of ants (Narendra, 2007; Philippides et al., 2011; Collett et al., 2013), where routes can be established by storing snapshots along a path to be navigated along. The snapshots can be stored either as they are (‘Perfect Memory’) or by feeding them into a holistic memory creation mechanism, such as the machine learning algorithm ‘Infomax’. The Infomax algorithm creates one holistic memory from all snapshots, memorizing effectively the most different snapshots from the whole set of snapshots (Baddeley et al., 2012; Wystrach et al., 2013).

Algorithms that rely on whole-body rotation or mental rotation (Möller, 2012) to evaluate possible directions of travel can be computationally or time intensive. Therefore, it is interesting to look in detail at the active scanning behaviours of insects to make snapshot navigation even more efficient. Based on the observation that ants move in sinusoidal trajectories, Kodzhabashev & Mangan replaced the stop-scan-go movement strategy with a sinusoidal movement pattern, where the amplitude is modulated by the snapshot familiarity. This leads to a high amplitude sinusoidal movement with weak input familiarity and a shallow sinusoidal movement with strong familiarity, enabling a robot to navigate along a route (Kodzhabashev & Mangan, 2015). Therefore, the scanning movement is spread out over several movement steps and makes the familiarity following process more dynamic.

In insects, rhythmic motor patterns are produced by a specific brain area called the Lateral Accessory Lobe. The LAL is a conserved brain structure, which receives inputs from learning and sensory centres and connects with the motor centres. It consists of two lobes, one located in each brain hemisphere. The inputs are largely divided hemispherically, meaning that the inputs towards one lobe mainly originate in the same hemisphere (Namiki et al., 2014) and subsequent processing occurs predominantly within the same hemisphere also (Paulk et al., 2015). Behaviourally, the LAL seems to be involved in the generation of small scale search behaviours (Kanzaki et al., 1992; Pansopha et al., 2014). These seem to be modulated by the reliability of navigation cues, leading to extensive search for a cue if that cue is weakly perceived, and targeted steering behaviours when the cue is strong (Namiki & Kanzaki, 2016; Steinbeck et al., 2020). Steering is achieved by an imbalance of activity of descending neurons towards the motor centres (Zorović & Hedwig, 2013; Iwano et al., 2010; Rayshubskiy et al., 2020). Because this appears to be a general purpose circuit in insects, we have recently developed a steering framework based on the LAL (Steinbeck et al., 2020), which provides a general mechanism for sensory information to control orientation and active search.

Based on the steering framework, where hemispherically divided inputs ultimately lead to a steering response, we investigate how bilaterally organised visual memories could achieve a steering response to navigate along a learned route. Instead of using a single panoramic image as a memory, as for many of the models above, we will use two lateralised memories. This is biologically plausible given the lateralised wide field eyes and the bilateral Mushroom Bodies, which are the site of visual route memories (Buehlmann et al., 2020; Kamhi et al., 2020; Li et al., 2020).

Given the LAL controls steering by using the reliability of lateralised sensory inputs, we ask how the organisation of bilateral view-based memories can be be optimised to provide a reliable steering signal. The investigation mainly focuses on the Field of View (FoV), which describes how much raw visual information is available to each hemispheric memory, and an Offset, which describes the target/familiar orientation of the stored memories in relation to the agent’s heading (Wystrach et al., 2020). The spontaneous difference in visual familiarity between the current and stored views for both hemispheres is used as a steering signal, similar to a taxis mechanism. In sum, we explore which type of visual system would fit best with this steering process and therefore would be a biological plausible input to the insect LAL during navigation.

## Models and methods

### Implementation

Modelling and analyses were performed with MatLab 2017 & 2019 (The MathWorks, Inc., Natick, Massachusetts, USA). The code to run the simulations and analysis as described in the methods can be found online at https://github.com/FabianSteinbeck/FamiliarityTaxis.

### Antworld Image Rendering

For the exploration of Bilateral Snapshot Navigation we chose to use the “Antworld” virtual environment, which is a LiDAR (Light Detection And Ranging) 3-D reconstruction of an experimental location, near Seville Spain, which has been used for observations and experiments with *Cataglyphis velox* ants (Risse et al., 2018). Within this environment we chose two routes which closely resemble real routes taken by ant foragers navigating towards their nest (routes 3 and 12 from the Brains on Board [BoB] robotics GitHub, figure 1 A). The rendering of the test images is done with the BoB robotics Antworld rendering pipeline. To generate training views and displaced test locations, on each route we chose 31 equally spaced locations, where the heading angle faces directly towards the next location. These are the training locations for the snapshots. For testing, at each location we rendered five off-route locations normal to the route direction, both to the left and right (distances were 5, 10, 30, 100, 300 mm; figure 1 D). The panoramic images are rendered so that the centre pixels represent the heading direction, i.e. the view orientation oscillated about the overall route direction.

**Fig. 1.**
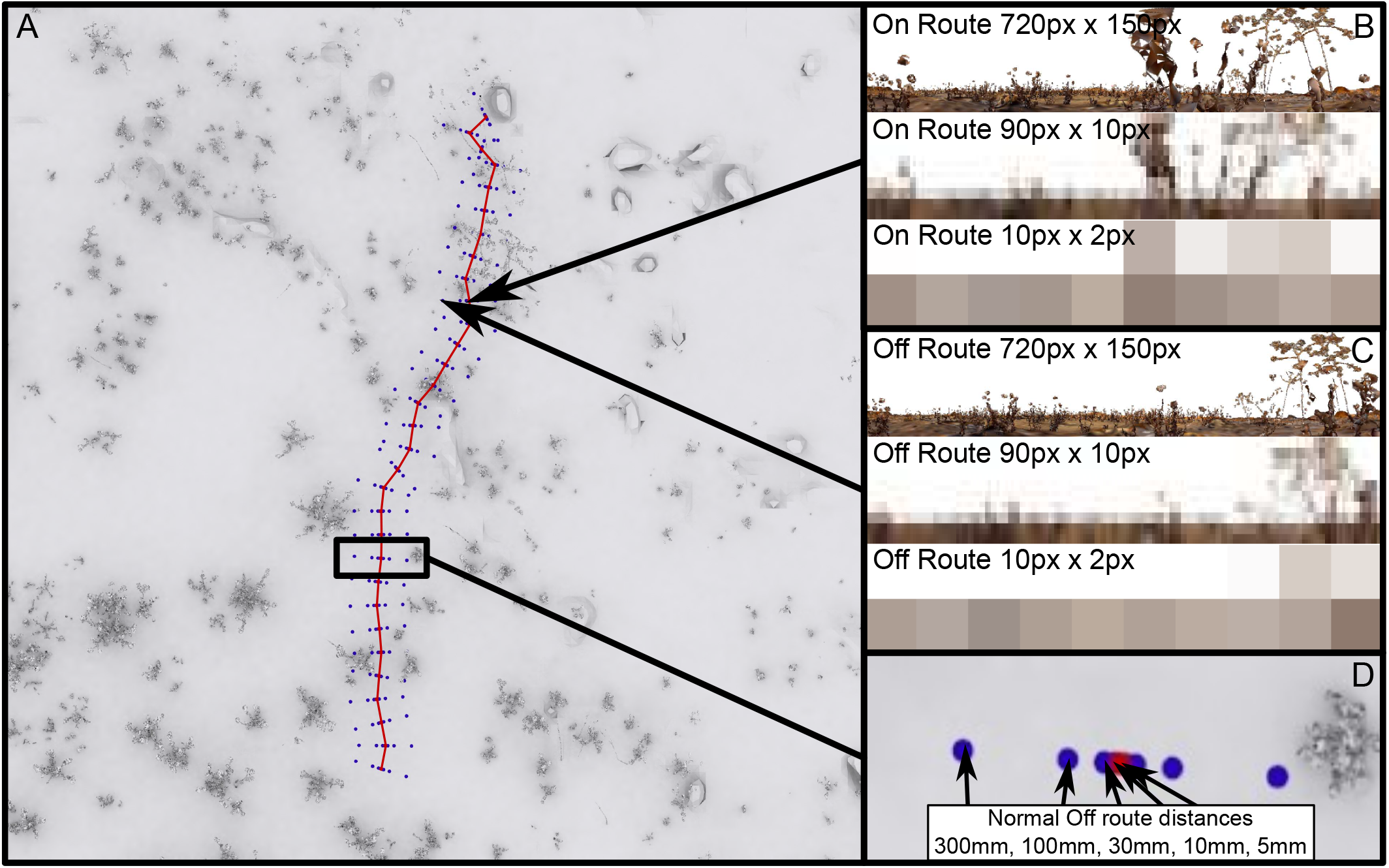
Visual environment “Antworld” simulating a desert-like environment with sparse vegetation. A top-down view of the virtual environment (A), showing Route A (red) and the off-route test locations (green) (D). The two locations singled out show the view from along the route (B) and displacements to the side of the route (C) at different visual resolutions.

### Field of View

The Field of View (FoV) describes the visual input available to each hemisphere/eye, which is then both used for snapshots and the comparison of the query images to the snapshots to generate the rotary Image Familiarity Function (rIFF). Because both hemispheres/eyes have a field of view, visual systems are further described by the degree of visual overlap (the visual range of both eyes overlapping in the front) and the size of the rear blindspot (how much visual range is missing to the back). A panorama’s central horizontal pixel is the heading direction, therefore the blindspot refers to how many pixels are taken away from the sides, and the overlap defines how many pixels are designated to be taken beyond the horizontal centre contralaterally for each of the images.

### Offset

The Offset describes the orientation of a stored snapshot in relation to an agent’s body and to the overall route direction. If a view is stored with no Offset this would mean that the most familiar direction is directly aligned with the agent’s overall direction of travel. An Offset angle for each FoV would mean that agents memorised a view when the body was rotated away by the Offset angle relative to the overall trajectory direction (figure 2, C). This would result in maximally familiar directions occurring at angles symmetric to the agent’s centre line. For example, an Offset of 45° would result in the left FoV having a maximum familiarity when the agent was rotated by 45° to the right from the original heading direction and the overall route direction from the training images, and vice versa (figure 2, C). For convenience within the setup of this experiment, this is achieved by rotating the panoramic query image for each hemispheric query image before cutting it into the FoV. The Offset values tested were [0°, 9°, 18°, 27°, 36°, 45°, 54°, 63°, 72°, 81°, 90°].

**Fig. 2.**
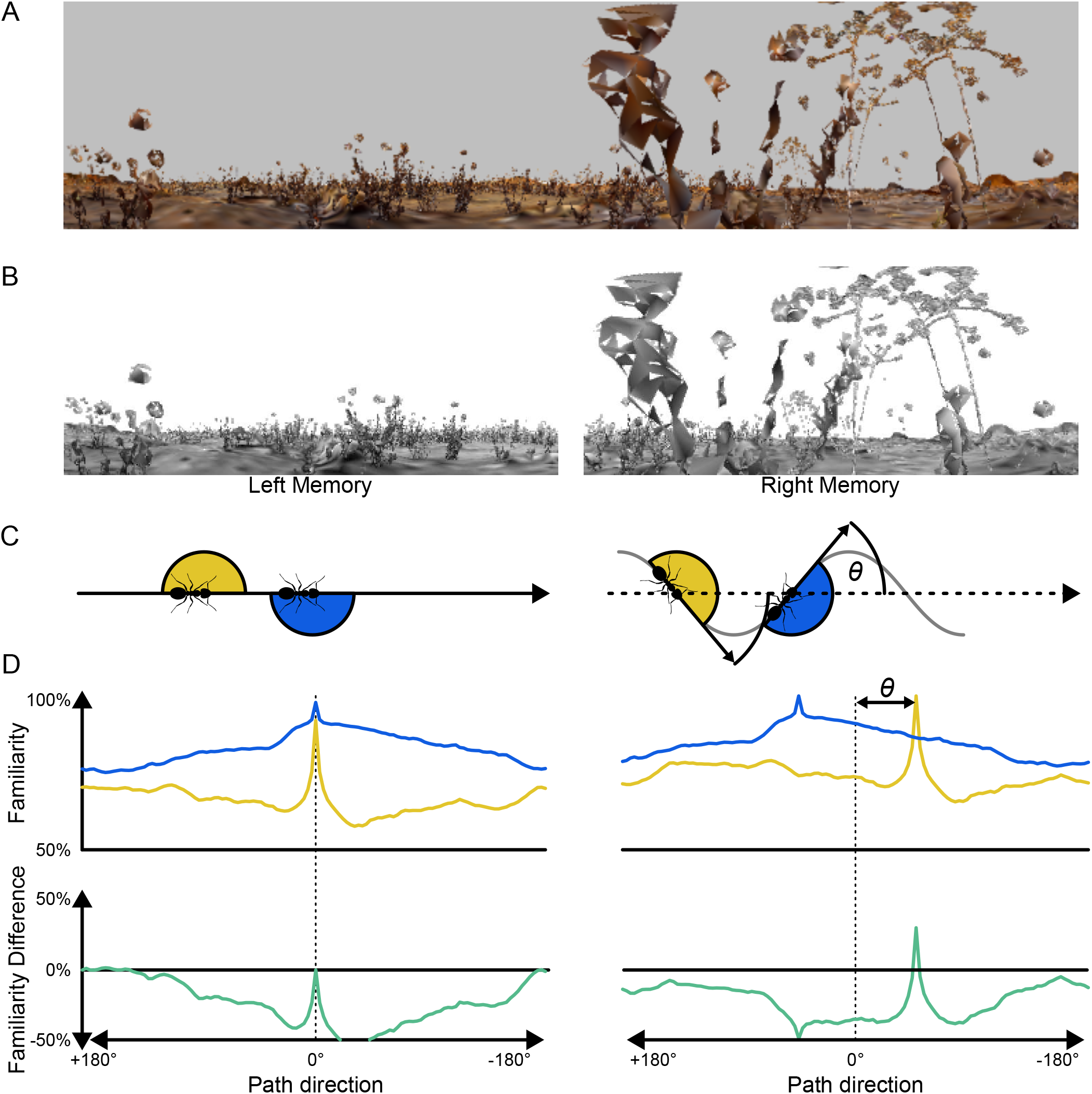
Impact of Offset on visual familiarity. A: A panoramic snapshot. B: The panoramic snapshot divided into two memories. C, left: Representation of when snapshots are taken with zero offset, i.e. when facing route direction; C, right: Snapshots taken with an Offset *θ*. D, left, top: rotary image familiarity functions per eye (yellow = left eye, blue = right eye) generated with the above snapshot/memories with no Offset. D, left, bottom: the difference between left and right eye familiarity functions. D, right: the same depictions, but for snapshots with an Offset.

**Table 1.** Field of View configurations with frontal Overlap and rear Blindspot values.

| FoV configuration | Overlap [°] | Blindspot [°] |
| --- | --- | --- |
| 1 | -180 | 0 |
| 2 | -90 | 90 |
| 3 | 0 | 180 |
| 4 | -90 | 0 |
| 5 | 0 | 90 |
| 6 | 90 | 180 |
| 7 | 0 | 0 |
| 8 | 90 | 90 |
| 9 | 180 | 180 |
| 10 | 90 | 0 |
| 11 | 180 | 90 |
| 12 | 180 | 0 |
| 13 | 270 | 90 |
| 14 | 270 | 0 |
| 15 | 360 | 0 |

### Image Processing

The rendered panoramic image dimensions are 720×150 pixels. First, the Sky is whitened and then the whole image converted to gray-scale. Each image then is cut into two views, depending on the specific FoV for that test. For investigations on the impact of resolution, images are downsampled to 90×10, 60×8, 30×6, 20×4 and 10×2 resolutions using MatLab’s imresize-function. For the main analysis we use 90×10, which gives a resolution comparable to the visual resolution of the ant compound eye (Schwarz et al., 2011).

### Rotary Image Familiarity Function

The Rotary Image Familiarity Function (rIFF) is used to systematically compare a stored snapshot with a series of rotated versions of the ‘current’ query image in order to find which orientation is the most familiar, i.e. the best match. The query image is rotated in steps through 360°, at each rotation step (3° for the 720 px image size, 1 pixel step for the smaller image sizes [4°, 6°, 12°, 18°, 36°]) the views for both eyes are extracted from the full panorama and then, with the given FoV, the covariance (familiarity) of each eye’s query image is calculated for the appropriate stored snapshot. Plotting the covariance for the whole 360° gives the rIFF, one for each hemisphere (figure 2 D).

### Measures of successful steering

Here, the displaced off-route images are compared to the on-route images at the nearest location. At each location, the rIFFs of the left hemisphere and right hemisphere are normalised to the *max −* 105% *· min* percentile before being subtracted from each other. This gives us a familiarity difference between left and right eyes which can be used to drive a steering response. Positive values suggest a leftwards turn and negative values indicate steering rightwards, examples where the familiarity difference gives a correct steering response are given in figure 2 bottom.

In order to assess the overall success of a given set of visual system parameters, we take the proportion of orientations, across all test locations, where the Left-Right difference in visual familiarity would indicate a correct turn response. We initially look at this for all orientations, i.e. for *±*180° from the ‘correct’ direction. However, the main focus of this paper is to consider how a bilateral familiarity metric can work for route guidance, in the context of the insect LAL brain region which modulates motor behaviour between steering (when sensory signals are strong) and search (when signals are weak). Therefore, for much of our analysis we focus on the proportion of orientations at test locations that give correct steering responses within *±*90° of the overall ‘training’ route direction. This gives us an appreciation of how likely the bilateral familiarity information would be in keeping an agent on the desired route, rather than relying on a search behaviour (see introduction).

### Evaluation of aliasing

When asking how bilateral visual familiarity measures might provide information to drive steering, we look at the likelihood of deriving the correct steering response when the views from displaced locations are compared to the nearest training/stored view from the target route. Real world agents might not have access to information that could ensure they compare their current view to the ‘correct’ training view. To analyse how aliasing (mistakenly selecting the wrong training view for comparison) might impact on overall performance, we looked at how rate of aliasing might vary for different visual parameters. The aliasing metric is derived by considering which of the stored training views gives the best match for the current location across all orientations and all possible stored views. This gives us a number for the difference in sequence location between the true best reference location and the position of the apparent best visual match. A score of 2 would mean that the best match was 2 positions away from the correct stored image location in the training route sequence of views.

## Results

### Overall performance of a bilateral visual familiarity method

Our primary goal for this analysis was to give proof of concept that bilateral visual familiarity information can be used to generate a steering signal (Steinbeck et al., 2020). The familiarity signal depends on the comparison of current views to stored route memories. The hypothetical scenario is that an agent has stored left and right eye views from along a foraging route and then when trying to navigate using those views, steering will depend on the comparison of left and right visual familiarity. That is, how similar are the current left and right eye views to the stored left and right views. The agent should steer towards the most familiar view, like a taxis mechanism.

In our first analysis, for all orientations at each test location, we evaluate whether the difference in the familiarity between the left and right visual field would lead to a correct steering response. Each pixel of the comparison plot between all visual systems represents one visual configuration (FoV, xaxis) and Offset (y-axis) combination, where we aggregate the tests from one route using all on-route and displaced test locations at all orientations. We then show the proportion of these tests (n = [31 on-route locations + 31 *·* 10 off-route locations] *·* 180°/3° rotations = 20,460) where the familiarity difference would give the correct steering response. This performance is only just better than chance when taking into consideration all possible orientations that an agent may find themselves (*±*180° relative to the heading direction), with 96% getting above 50% correct, but only 1/165 getting 75% correct. However, our target behaviour is visual route following we are only taking into consideration the directions *±*90° relative to the heading direction, 51/165 (31%) get at least 75% correct and 10/165 (6%) get at least 80% correct. Thus we have existence proof that there are a range of visual configurations that can generate accurate bilateral familiarity based steering for view based navigation along a route.

### Organisation of visual system

The secondary goal of the work was to investigate how performance depends on the precise organisation of the bilateral visual field sizes and positions, as well as the orientation (offset) of the left and right visual memories.

The configurations of FoV and Offset resulting in at least 75% of correct steering responses are found amongst the larger FoVs and non-zero Offsets, 43 out of 165 (26%) visual system configurations at 720×150 resolution performed over 75% (figure 3A, cyan). 80% is reached only with the largest FoVs and medium to large Offsets (figure 3A, magenta). When analysing the influence of the Offset in more detail, it becomes apparent, that any Offset other than zero leads to a much improved performance and performance quickly plateaus across a wide range of Offsets (figure 3B). The values fall off towards 90° since the range of considered orientations is *±*90°, and for the larger Offsets, correct steering responses may occur beyond this threshold. Investigating in detail how the correct steering responses are distributed for one visual configuration and different Offsets, shows how correct steering responses cluster around the Offset angles (figure 3C C). That means that the Offset angle plays a crucial role in generating a correct steering response using the difference of familiarity between the left and right view comparisons.

**Fig. 3.**
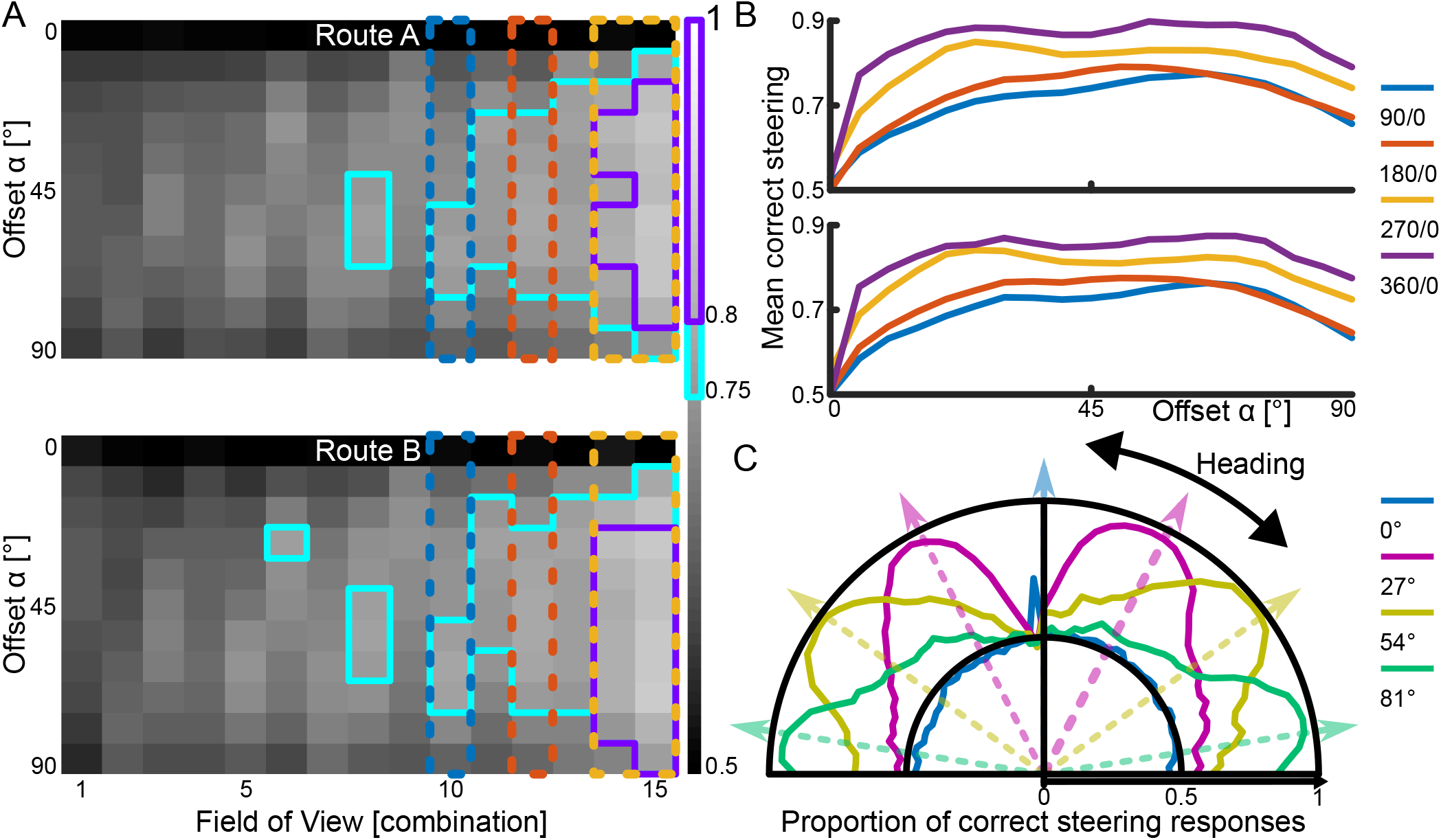
Steering performance for visual system configurations. A: Each square represents the proportion of correct steering responses over all tested locations and for orientations within 90° of the overall route direction for a particular visual configuration (x-axis) and Offset (y-axis). The cyan framed pixels surpass 75% and the magenta framed pixels surpass 80% correct steering responses. The colourful columns highlight the visual configurations used for B. B: Both top and bottom show performance relative to more granular resolution of Offsets for routes A and B respectively. C: Proportion of correct steering responses for each orientation, using a visual configuration with 90° Overlap and 90°Blindspot as well as a range of Offsets indicated by the line colours.

### Visual resolution

Reducing the visual resolution increases the proportion of correct steering responses. 45 out of 165 (27%) visual system configurations performed over 75% with 90×10 resolution, 53 out of 165 (32%) with 60×8 resolution, 56 out of 135 (41%) with 30×6 resolution, 45 out of 90 (50%) with 20×4 resolution, 34 out of 60 (57%) with 10×2 resolution. The analyses was performed on two routes, but the pattern of results was very similar, therefore were combined them (figure 4A). The increase in performance with reduced resolution matches previous findings regarding using different visual configurations for snapshot navigation (Milford, 2013; Wystrach et al., 2016).

**Fig. 4.**
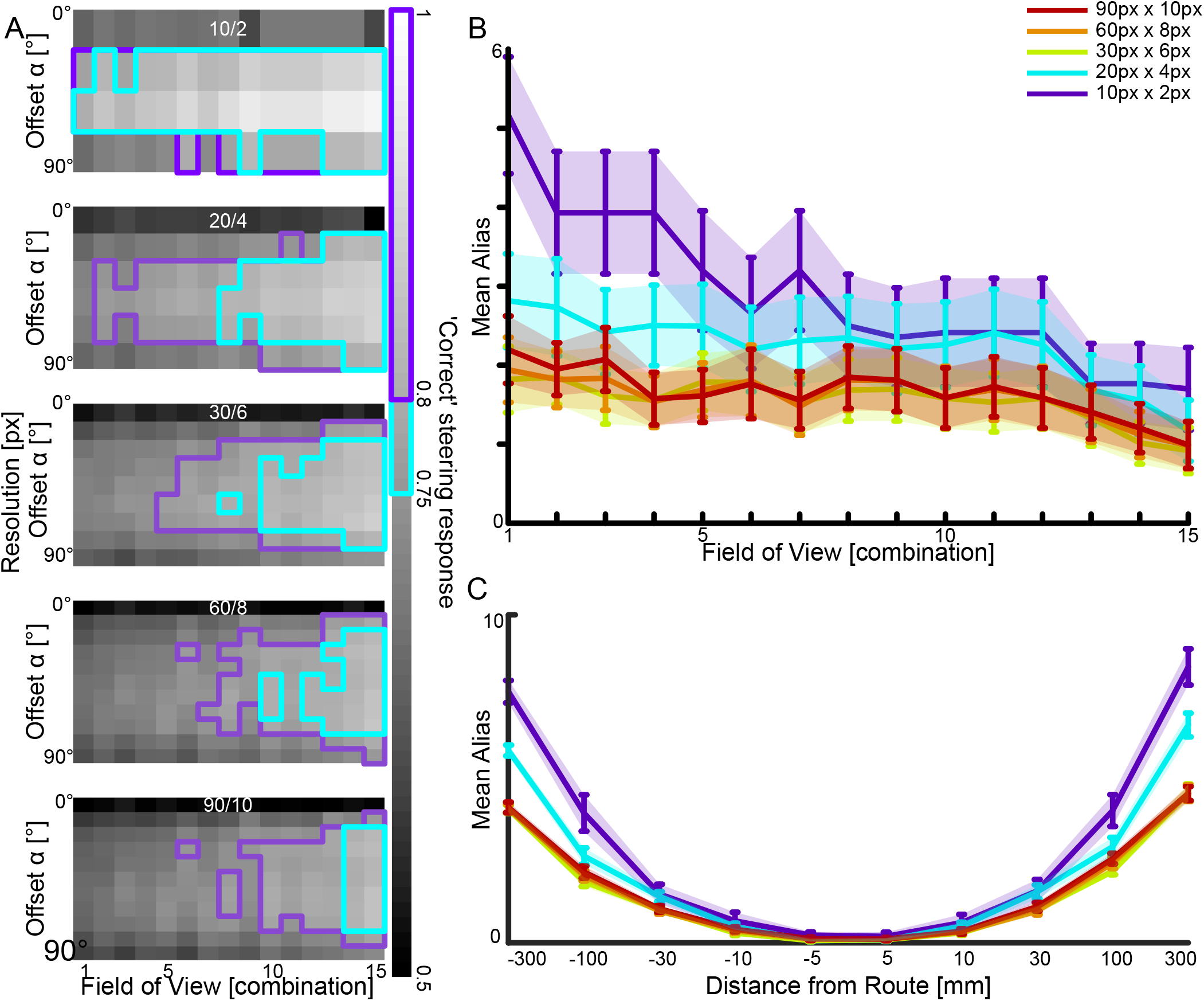
The impact of visual resolution and aliasing. A: The correct steering performances for the visual system configurations with different visual resolutions, when using the correct memories. The resolution is shown on top of each image (horizontal resolution/vertical resolution). The cyan framed pixels surpass 75% and the magenta framed pixels surpass 80% correct steering responses. B: For each visual system configuration we calculated the mean aliasing along the routes, with different visual resolutions. C: We show the mean aliasing relating to the distance from the route for different visual resolutions.

### Aliasing

For our primary analyses we calculated the familiarity of left and right views using the stored route view that was closest to the test location, i.e. the correct view. However, in natural scenarios, when ants are displaced from the route, the best matching view may be a view from an incorrect location at another place along the route, so called aliasing. We investigated how aliasing depended on visual specification and resolution. Aliasing occurs for all tested configurations, but less so for higher resolutions and larger FoVs (figure 4B). Aliasing also increases with displacement distance from the training route (figure 4C). When aliasing is introduced to testing the correct steering responses to Route A, the correct steering performance across all visual system configurations reduces to 94% compared to the perfect memory performance. This means, instead of comparing the query view to the correct memory in the memory sequence, all memories are evaluated, the highest familiarity match chosen which would be an incorrect view in 6% of cases.

## Discussion

### Takehome message

Steering in insects is achieved by an imbalance of activity between left and right descending neurons towards the motor centre. As sensory computation pathways are hemispherically divided, we investigated how two Fields of View instead of a single panoramic Field of View can aid in Snapshot navigation. To do this, we explored how a difference in familiarity between a left and right Field of View could achieve a steering response that would keep an agent on a route. We found that by subtracting the familiarity values from one hemisphere from those from the contralateral hemisphere we got at least 75% ‘correct’ steering responses towards the goal-direction from off-route locations within *±*90° of the route direction, given large Field of Views and medium to large Offsets. This performance increases with reduced visual resolution and is only slightly negatively affected by aliasing. Therefore, we have shown the plausibility of a steering mechanism based on the inter-hemispheric difference in image familiarity for snapshot navigation.

### Building on pixel-by-pixel comparison

Recent advances in insect neuroscience have inspired other ways in which view-based navigation models can be implemented efficiently. In regard of visual memory, the Mushroom Bodies (MB) have been identified as a structure capable of forming memories of visual stimuli (Liu et al., 1999; Vogt et al., 2014; Buehlmann et al., 2020; Kamhi et al., 2020; Li et al., 2020). A plausible model of how Mushroom Bodies might implement visual memory for navigation comes from Ardin et al. Visual projection neurons connect randomly to a 2nd layer, where each 2nd layer neuron needs activation of multiple inputs to fire. These converge then onto one extrinsic neuron, which introduces a reinforcement signal, therefore associating a certain combination of inputs to a reward. This leads to a measure of familiarity. The number of neurons determine the capacity of this network, scaling the capacity logarithmically. Using this type of memory in a Snapshot model generates comparable performance to other algorithms (Ardin et al., 2016).

For processing of the images we used pixel-by-pixel comparisons, this stage of processing was not intended to be biologically plausible, which made the familiarity processing a simple mathematical operation. This approach can lead to several hurdles though, especially in dynamically changing environments, for example, the same location’s raw pixel value representation may change dramatically with different lighting. To compensate for this, several image processing methods have been proposed. While some methods are similar to feature detection, as with skyline (Graham & Cheng, 2009), landmark (Möller et al., 1999) or Haar-like features (Baddeley et al., 2011), others take a holistic frequency filtering approach (Stone et al., 2018; Meyer et al., 2020). More computationally intensive methods like object recognition could be employed, albeit probably for simple shapes given low resolution insect vision. Regardless, our approach can be seen to be a minimal demonstration of the recovery of an accurate steering response, any additional ‘cognitive’ processes will therefore increase the performance and/or make it more robust to environmental changes. Another filtering approach, which we did investigate, is downsampling of the image size, which effectively acts as a low-pass filter, thereby cancelling out high-frequency noise, as is inherent in the low resolution visual systems of insects (Stürzl & Mallot, 2006; Gerstmayr et al., 2008; Baddeley et al., 2011). In our analyses, downsampling had a positive effect on performance, as has been shown for visual navigation tasks previously. (Milford, 2013; Wystrach et al., 2016).

In our modelling, visual configurations allowed the independent left and right views, to have a field of view extending into the contralateral side, i.e. a visual overlap. The extent to which visual information is transferred to the contralateral hemisphere is currently unclear (Habenstein et al., 2020; Li et al., 2020). However, indicated by our best performing visual setups, the more visual information each memory can capture, the better performance. However, it is also true that a FoV using only ipsilateral visual information can be successful too, if in combination with a medium Offset (for example FoV 90°/90°. see figure 3). A previous investigation of multiple visual fields for snapshot navigation yielded increased performance with increasing number of visual fields, with two fields being a plausible trade-off between the number of visual fields of view and the computational effort of determining rIFFs (Wystrach et al., 2016).

### What to remember

While we mainly focused on investigating how the hemispheric familiarity difference mechanism works when the views from off route locations are compared to the ‘correct’ memory, the matching process in an autonomous agent may be less accurate and aliasing may occur, where the most similar snapshot actually originates from a location at another route location (Knight et al., 2020). Aliasing typically reduces performance and comparing each query image to all stored route snapshots can become computationally expensive, as this method scales linearly with the snapshot number. One attempt to reduce aliasing is inspired by ants’ apparent sequential memory, where they expect certain views to occur one after another (Schwarz et al., 2020). Algorithmically, a temporal window can be introduced, where only a limited amount of the sequentially stored snapshots is used for familiarity detection. This way, general aliasing can be minimised as well as more complex routes followed (Kagioulis et al., 2021).

Additional robustness of view based navigation might also be gained from different organisations of view memories. For instance, one could be the distinction between attractive and repulsive memories, where the agent would not only take snapshots of the direction to move towards (attractive, towards the goal), but also of the direction not to move towards (repulsive, 180° away from the goal, Le Möel & Wystrach (2020)). This process resulted in a fine scale sinusoidal movement trajectory (oscillations). While we used an attractive memory, an additional repulsive memory could emphasize a steering response with the attractive memory pulling towards the goal-direction and the repulsive memory pushing away from the anti-goal-direction. Yet another approach could be used to tie the panoramic memories to steering instructions. Whenever heading direction is to the side of the overall goal direction (similar to our offset), that view could be tagged with the appropriate steering direction, where the steering instruction could originate in the LAL (Wystrach et al., 2020).

Similar to Wystrach et al. (2020) we also found that an Offset increases a model’s success in steering. This would indicate that snapshots would have to be captured not when an agent’s heading is aligned to route direction, but when the agent’s heading is at an angle. Many ant species oscillate along the route they are following (Graham & Collett, 2002; Clement et al., 2022) which could be part of the strategy of storing the snapshots with an Offset. We found that best steering signal are to be found near the Offset of the stored views (figure 3C). Thus for our method and for Wystrach et al. (2020) oscillations when navigating would come from the agent ‘bouncing’ back and forth between leftward and rightward steering. Since the observed oscillations in ants are rather regular however, this could also mean that a source of the oscillations may be internally generated (Clement et al., 2022), perhaps from the LAL (Steinbeck et al., 2020).

### Central place navigation task

Insect neuroscience has provided us with a detailed understanding of the brain areas important for orientation and spatial control, the Central Complex (CX). It is a conserved brain structure which receives multimodal inputs (Heinze et al., 2013). It is involved in spatial orientation such as compass driven behaviours (Cope et al., 2017; Pisokas et al., 2020; Dan et al., 2021) and may also play a role in view-based navigation in ants (Wystrach et al., 2020). The CX receives Mushroom Body outputs, and might be used to maintain a course that has been set by visual information (Schwarz et al., 2017).

To demonstrate the potential of a bilateral familiarity in a biologically plausible neural model of motor control. We integrated bilateral familiarity information with a LAL model (Steinbeck et al., 2022) for a relatable navigation task, which is navigating towards a central place (see figure 5). For this, we took six snapshots around a central place, each one heading towards the central place, at three distances (0.5m, 1m, 1.5m), therefore 18 in total. This resembles the images that might be stored following learning walks in ants, where they circle around the nest and periodically look back at it, thereby potentially memorizing views of the nest from different directions and distances(Philippides et al., 2013). We generated a grid around that central place spanning 8 metres into the horizontal plane with a resolution of 0.1 metres. At each grid location, we calculated the familiarity of each eye individually given the current view and the stored snapshots. The familiarity values were fed into a LAL-SNN, as described in Steinbeck et al. (2022). The LAL-SNN is a sensory stimulus pursuing network inspired by the insect brain with two main characteristics. The first is the usage of sensory inputs which are designated by brain hemisphere origin and relayed by the CX, as in all left hemisphere processed stimuli are fed into the left half of the network and vice versa. The second aspect is the formation of a Central Pattern Generator (CPG), which generates phasic switches of activity depending on the input strength between the two halves of the network. In effect, if a navigation relevant stimulus is strongly perceived, the network generates a deterministic steering behaviour towards that stimulus; if the perception of the focal stimulus is weak, it generates small scale search behaviours, therefore actively searching for relevant stimuli. Such a described agent was spawned in each grid location, clearly the agents follow routes and navigate in many cases to the central place (figure 5). Whilst this simulation is not meant to be 100% biologically, and real world, realistic, it does given existence proof that bilateral familiarity fed into LAL inspired SNN model can drive navigational behaviour.

**Fig. 5.**
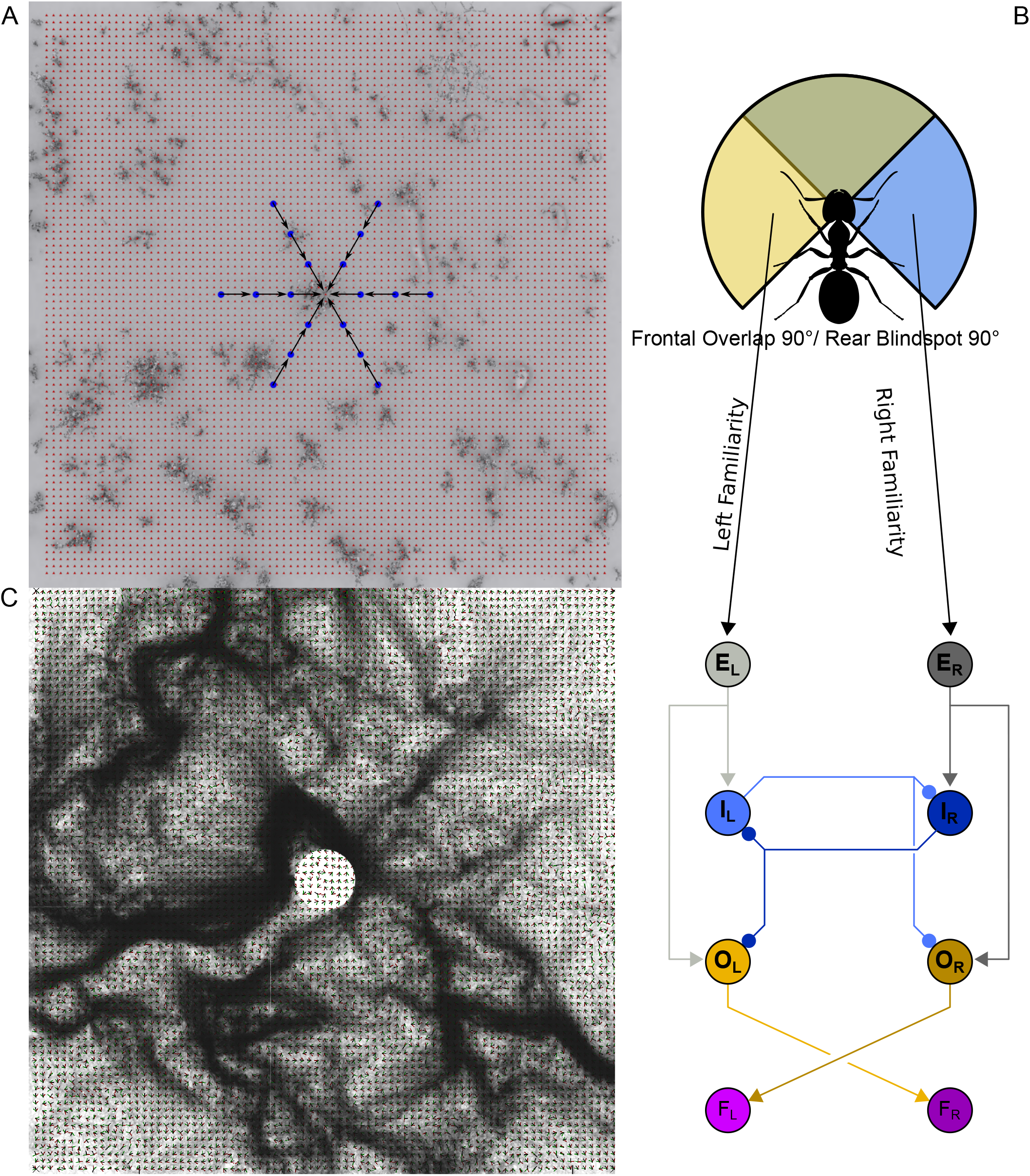
A central place navigation task. A: Memories are taken, aiming at a central place (blue dots). Around the area, a grid of snapshots (red dots) is set up, which are used to pre-calculate familiarity using these memories. B: The agent uses a visual system configuration of 90° Overlap and 90° Blindspot. The familiarity values are drawn from the closest grid point, and fed into a Spiking Neural Network (Steinbeck et al., 2022). C: Paths from agents released from each grid point where their steering comes from the SNN driven by familiarity information.

## Conclusion

Snapshot navigation has long been investigated using monocular panoramic views. This approach has been shown to work well in many scenarios. However, recent studies suggest that each ant eye’s view matters on its own. Both disabling the view of one eye affects the navigational capabilities (Buehlmann, unpublished data, Schwarz et al. (2023)) as does disabling the memory substrate (MB) of one hemi-sphere (Buehlmann et al., 2020). As our study suggests a familiarity relationship between both hemispheres could be used to recover the goal direction we hope that the idea of hemispheric imbalance will spark similar kinds of modelling attempts within computational neuroscience as well as further behavioural experiments.

